# Long-lasting reduction of intrinsic excitability in O-LM interneurons is mediated by endocannabinoid-dependent up-regulation of Kv7 channels

**DOI:** 10.1101/2021.01.08.426009

**Authors:** Salvatore Incontro, Malika Sammari, Norbert Ankri, Michael Russier, Jacques Fantini, Dominique Debanne

## Abstract

KCNQ-Kv7 channels are found at the axon initial segment of pyramidal neurons where they control cell firing and membrane potential. In oriens lacunosum moleculare (O-LM) interneurons, these channels are mainly expressed in the dendrites, suggesting a peculiar function of Kv7 channels in these neurons. The physiology of Kv7 channels is well characterized today but the precise contribution of these channels to neuronal plasticity is still unknown. Here, we show that Kv7 channel activity is up-regulated following induction of presynaptic long-term synaptic depression (LTD) in O-LM interneurons, thus resulting in a synergistic long-term depression of intrinsic neuronal excitability (LTD-IE). Both LTD and LTD-IE involve endocannabinoid (eCB) biosynthesis for their induction. Molecular modeling shows strong interaction of eCBs with Kv7.2/3 channel, suggesting a persistent action of these lipids on Kv7 channel activity. Our data thus unveil a major role for eCB synthesis in triggering both synaptic and intrinsic depression in O-LM interneurons.

## INTRODUCTION

Interneurons are fundamental components of the cortical area, orchestrating the complex machinery that governs brain rhythms. The control of pyramidal cell activity by GABAergic interneurons is required for the execution of hippocampal functions during spatial memory formation. Hippocampal interneurons consist of a very diverse population of cell types that have distinct postsynaptic domains and therefore differentially control input/output activity (1). In the hippocampus, parvalbumin (PV) and somatostatin (SOM) expressing interneurons form two broad subtypes of interneurons that preferentially target perisomatic and distal dendritic regions of pyramidal neurons, respectively, and are active on different phases of the theta cycle (2–4).

Oriens lacunosum moleculare (O-LM) interneurons are SOM-positive and selectively active during theta oscillations, gamma oscillations and fast ripples (5). They produce feed-back inhibition to pyramidal neurons. Receiving glutamatergic inputs from pyramidal neurons they not only inhibit the apical dendrites of CA1 pyramidal neurons in the lacunosum moleculare region but they also inhibit interneurons located in the stratum radiatum (6). Therefore, they may differentially modulate glutamatergic inputs from CA3 neurons and from entorhinal cortex respectively via the control of inhibition of the dendritic region where Schaffer collaterals project or via direct inhibition of the dendritic region where temporoammonic pathway project. They receive glutamatergic inputs from CA1 pyramidal neurons and cholinergic inputs from the medial septum (6). O-LM interneurons express a wide range of voltage-gated channels including Kv7.2/3 channels in their dendrites (7).

KCNQ-Kv7 potassium channels are found at the axon initial segment of pyramidal neurons where they control cell firing and resting membrane potential (8). Kv7 channels oppose depolarization by creating a non-inactivating outward current. These channels are inhibited by stimulating many receptors including muscarinic receptors (9). On the other hand, they are activated by somatostatin and by eCBs (10). While their redistribution along the axon initial segment has been demonstrated in homeostatic plasticity (11), their contribution to functional plasticity is still unclear.

Classically, activity-dependent plasticity of inhibitory circuits is thought to be achieved by regulation of excitatory synaptic drive to inhibitory interneurons (12–18) or by plasticity of inhibitory synaptic transmission on pyramidal neurons (19, 20). However, functional plasticity might also be achieved through the regulation of voltage-gated ion channels that control synaptic integration and spike initiation (21). PV interneurons express long-lasting intrinsic plasticity following induction of long-term synaptic potentiation (LTP) or activity deprivation (22, 23). However, it is still not known whether SOM interneurons as O-LM cells express intrinsic plasticity.

In hippocampal neurons, synaptic and intrinsic plasticity are expressed in parallel (21). This synergy is functionally important as both forms of plasticity act in concert to either promote or reduce excitation. However, the molecular mechanisms linking the two forms of plasticity are still unclear. Here, we show that long-term synaptic depression (LTD) and long-term depression of intrinsic neuronal excitability (LTD-IE) expressed in O-LM interneurons are both mediated by lipid biosynthesis. Stimulation of CB1 receptors induces LTD whereas the direct interaction of eCBs with Kv7.2/3 channels induces LTD-IE. Our results thus provide a previously unexpected involvement of eCBs in long-lasting plasticity of intrinsic excitability in GABAergic interneurons.

## RESULTS

### Co-induction of long-term synaptic and intrinsic depression in O-LM cells

Whole cell recordings from CA1 O-LM interneurons were obtained in hippocampal slices from P14-P16 rats. O-LM cells were identified in the *stratum oriens* by their typical electrophysiological behavior (regular spiking discharge with deep after-hyperpolarization and the presence of depolarizing sag upon hyperpolarizing current injection (24)) and their characteristic axonal projection in the *lacunosum moleculare* (**Fig. 1A**). All experiments were performed in the presence of the GABA receptor antagonist picrotoxin (PiTx, 100 μ M). Excitatory postsynaptic potentials (EPSPs) were evoked by stimulation of the axons of pyramidal neurons in the *alveus-oriens* region and excitability was monitored before and after pairing over a period of > 40 minutes (**Fig. 1B**). LTD was induced by pairing single action potential triggered in the O-LM cell with presynaptic stimulation delayed by 10 ms in groups of 6 at 10 Hz (18). As expected, a nearly 50% decrease in the EPSP slope was observed after such negative pairing protocol (52 ± 3% of the control EPSP slope, n = 16; **Fig. 1C**). An increase in the paired-pulse ratio was observed after induction of LTD (from 1.56 ± 0.18 to 2.83 ± 0.27, n = 5; Wilcoxon test, p<0.05; **Fig. S1A**), indicating that it is expressed by a presynaptic reduction in glutamate release.

**Figure 1.**
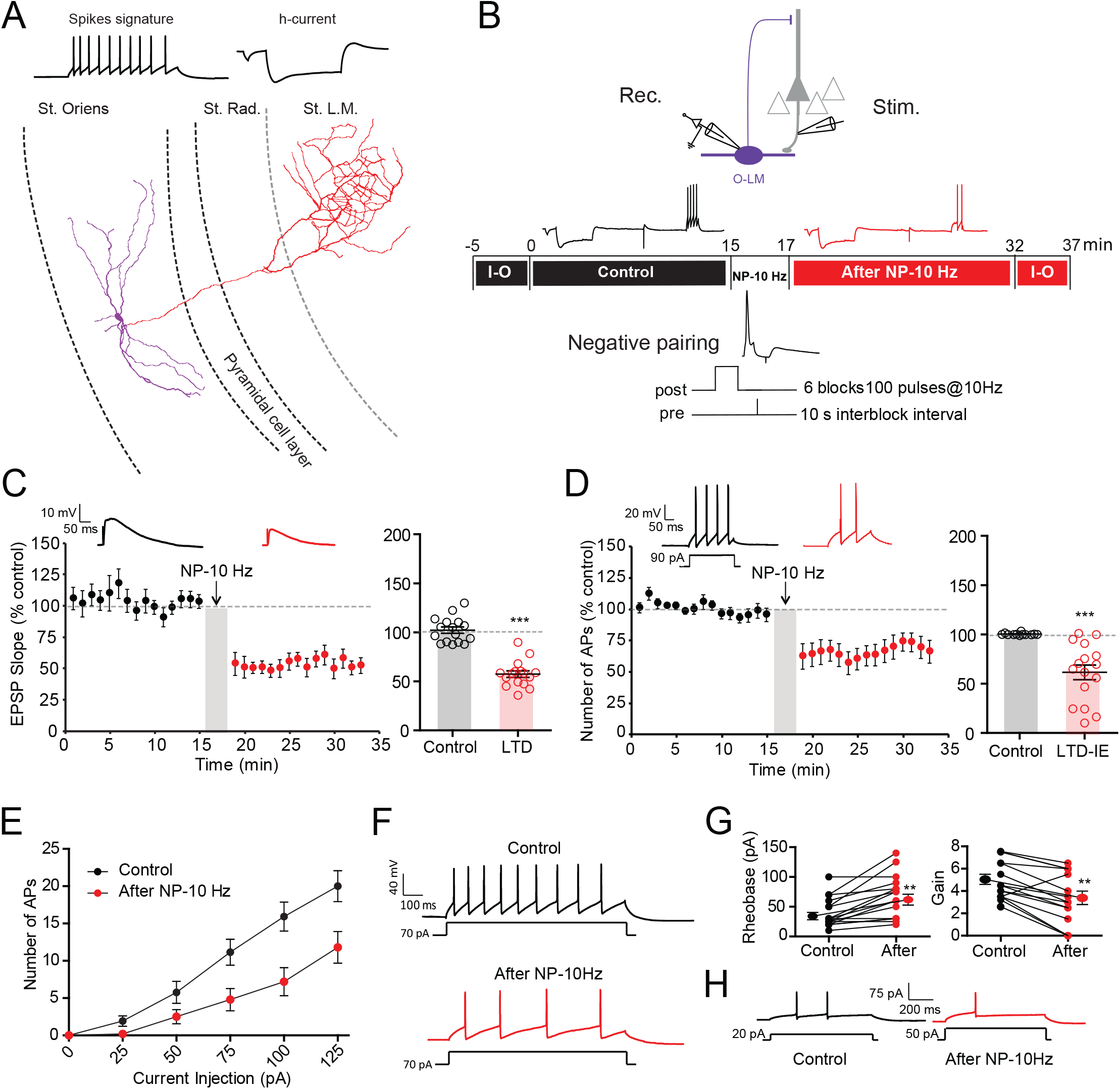
Negative pairing at 10 Hz induces LTD and LTD-IE in O-LM interneurons. A. Morphological and electrophysiological identification of O-LM interneurons. Reconstruction of an O-LM interneuron. Purple segments represent dendrites and red segments represent axon. B. Top, recording and stimulation configuration. Middle, synaptic transmission and intrinsic excitability is monitored at least 15 minutes before and after negative pairing delivered at 10 Hz (NP-10 Hz). Input-output (I-O) curves are plotted before and after the two test periods. Bottom, induction protocol. C. LTD induction by negative pairing. Left, time course. Right, pooled data. ***, Wilcoxson, p<0.001. D. Induction of LTD-IE by negative pairing. Left, time course. Right, pooled data. ***, Wilcoxson, p<0.001. E. Input-output curves before and after negative pairing. F. Reduced O-LM firing after negative pairing. G. Rheobase is elevated whereas gain is reduced. **, p<0.01. H. Example of the rheobase increase (here from 20 pA to 50 pA) after negative pairing.

Interestingly, the number of action potentials evoked by moderate current injections (90-120 pA, 100 ms) was significantly reduced after negative pairing (69 ± 6% of control AP number, n = 16; **Fig. 1D**). Importantly, no change in input resistance was observed (100 ± 3%, n = 16; **Fig. S1B**). Input-output curves drawn before and after pairing revealed a marked reduction in intrinsic excitability (**Fig. 1E** and **Fig. 1F**). The rheobase (i.e., the minimal current eliciting an action potential) was found to be elevated (from 40 ± 6 pA to 67 ± 9 pA, n = 16, Wilcoxson, p<0.01; **Fig. 1G**). This depression of intrinsic excitability could be followed up to 30 min (**Fig. S1C**), indicating that it is long lasting. Thus, negative pairing in O-LM interneurons leads to the co-induction of LTD and long-term depression of intrinsic excitability (LTD-IE). LTD and LTD-IE were found to be linearly related (**Fig. S1D**), suggesting a common induction mechanism.

### Induction mechanisms of LTD-IE in O-LM interneurons

As LTD and LTD-IE are induced by negatively pairing postsynaptic APs with EPSPs delayed by 10 ms at a frequency of 10 Hz, we checked the associative nature of the stimulation as well as the frequency dependence.

Post-synaptic spiking alone delivered at 10 Hz was unable to induce either LTD (101 ± 3% of control EPSP slope, n = 8) or LTD-IE (93 ± 4% of the control AP number, n = 8; **Fig. 2A-C**), whereas in a large fraction of the same neurons, a second episode of post-pre pairing induced both LTD (67 ± 8%, n = 6) and LTD-IE (70 ± 5%, n = 6; **Fig. 2A-C**). Similarly, pre-synaptic stimulation alone delivered at 10 Hz was unable to induce either LTD (106 ± 7%, n = 11) or LTD-IE (98 ± 6%, n = 11) whereas in a large fraction of the same neurons a second episode of post-pre pairing induced both LTD (71 ± 8%, n = 9) and LTD-IE (71 ± 8%, n = 9). These results show that LTD and LTD-IE induction in O-LM interneurons depends on the association between post-synaptic spiking and presynaptic stimulation.

**Figure 2.**
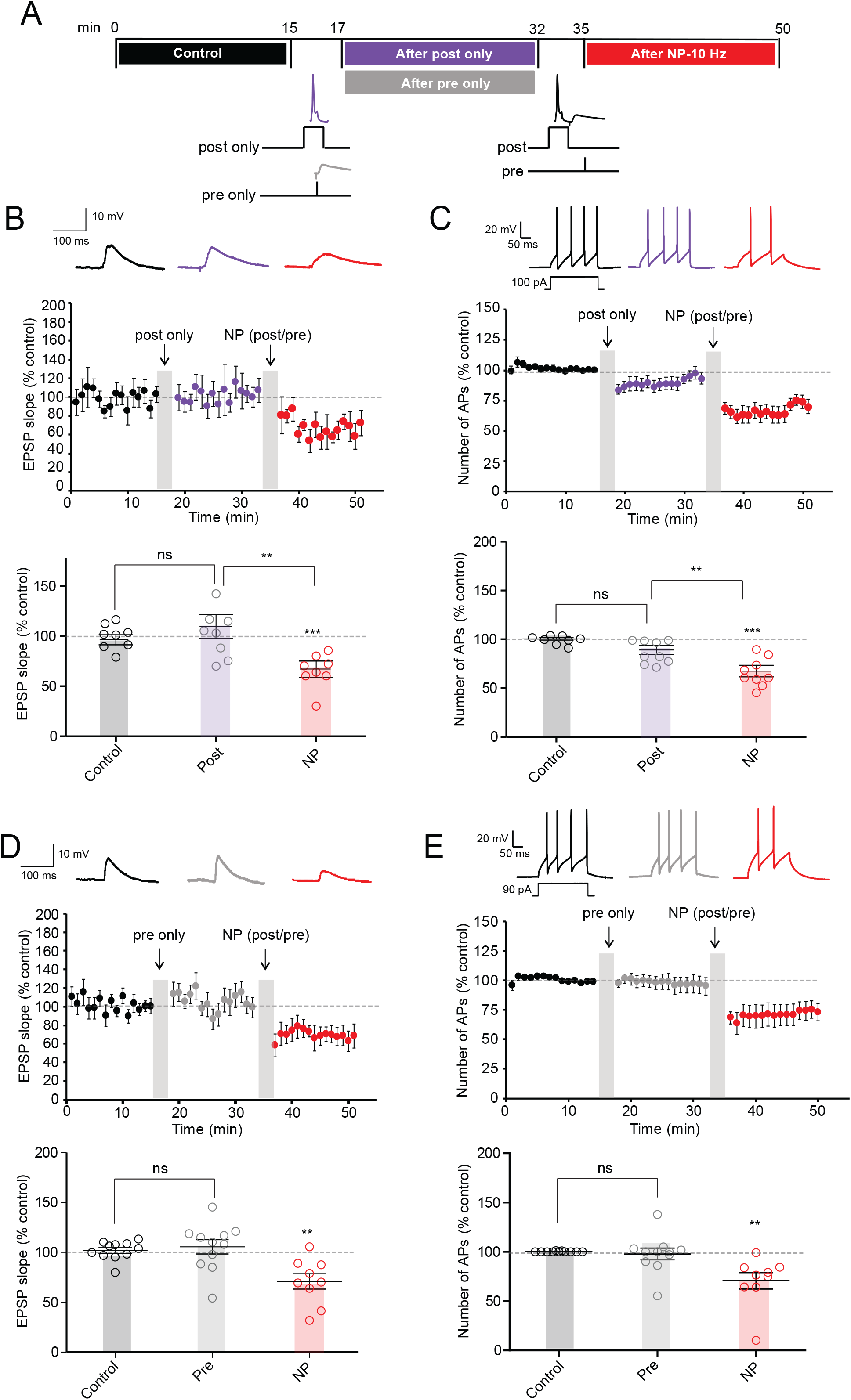
Post-pre stimulation is required for induction of both LTD and LTD-IE. A. Induction protocols. In a first step, the effects of post-synaptic or pre-synaptic activity (“post only” or “pre only”) is tested. Then, in each case, a post-pre protocol delivered at 10 Hz is applied. B & C. Post-synaptic spiking activity results in no synaptic or intrinsic changes whereas post-pre pairing in the same neurons induced LTD and LTD-IE. D & E. Pre-synaptic activity alone does not change synaptic transmission nor intrinsic excitability whereas post-pre pairing is sufficient to induce LTD and LTD-IE in the same neurons.

In a second step, we varied the stimulation frequency during pairing. At 5 Hz, no significant LTD (73 ± 11%, n = 10, Wilcoxson, p>0.1) nor LTD-IE (98 ± 8%, n = 10, Wilcoxson, p>0.1) was induced (**Fig. S2**). Similarly, no LTD (89 ± 16 %, n = 7; Wilcoxson, p>0.1) and a very small LTD-IE (81 ± 6, n = 7; Wilcoxson, p<0.01) could be induced by stimulation at 20 Hz (**Fig. S2**). Thus, we conclude that LTD and LTD-IE are preferentially induced by stimulation at a frequency of 10 Hz.

### LTD-IE in O-LM interneurons is mediated with Kv7 channels

We next focused on the expression mechanism of LTD-IE expressed in O-LM interneurons. As an increase in the rheobase was observed after LTD-IE induction, we first checked whether the voltage threshold of the action potential (AP) was modulated after negative pairing. However, no difference in the AP threshold was observed before and after induction of LTD-IE (−43.3 ± 1.4 mV in control vs. −42.4 ± 1.2 mV; Wilcoxson, p>0.1; **Fig. S3A**). This result thus excludes a change in Nav or Kv1 channels controlling the spike threshold. Moreover, no change in input resistance measured by long hyperpolarizing steps of current was observed, thus excluding a major role of HCN channels in LTD-IE expression.

As O-LM interneurons express a high density of Kv7 channels in their dendrites (7), we next focused on the Kv7 mediated M-type current. As HCN and Kv7 may have overlapping deactivation and activation profiles, we first measured the M-current before and after induction of LTD-IE in the presence of the specific blocker of HCN channels, ZD-7288 (1 μM; **Fig. S3B-D**). To measure the M-current, O-LM neurons were recorded in voltage clamp at a membrane potential of −30 mV and negative pulses to −50 mV were applied to measure the deactivation of M-current. Interestingly, the magnitude of the M-current was significantly increased following induction of LTD-IE (from 26.8 ± 7 pA to 54.2 ± 11.4 pA, n = 14; **Fig. 3A** and **3B**). To confirm that M-current was effectively measured, the specific antagonist of Kv7 channels, XE-991 (1 μM) was applied at the end of the recording in a few experiments. Most of the current was shown to be blocked in the presence of XE-991 (remaining current: 19.7 ± 4.9 pA, n = 8).

**Figure 3.**
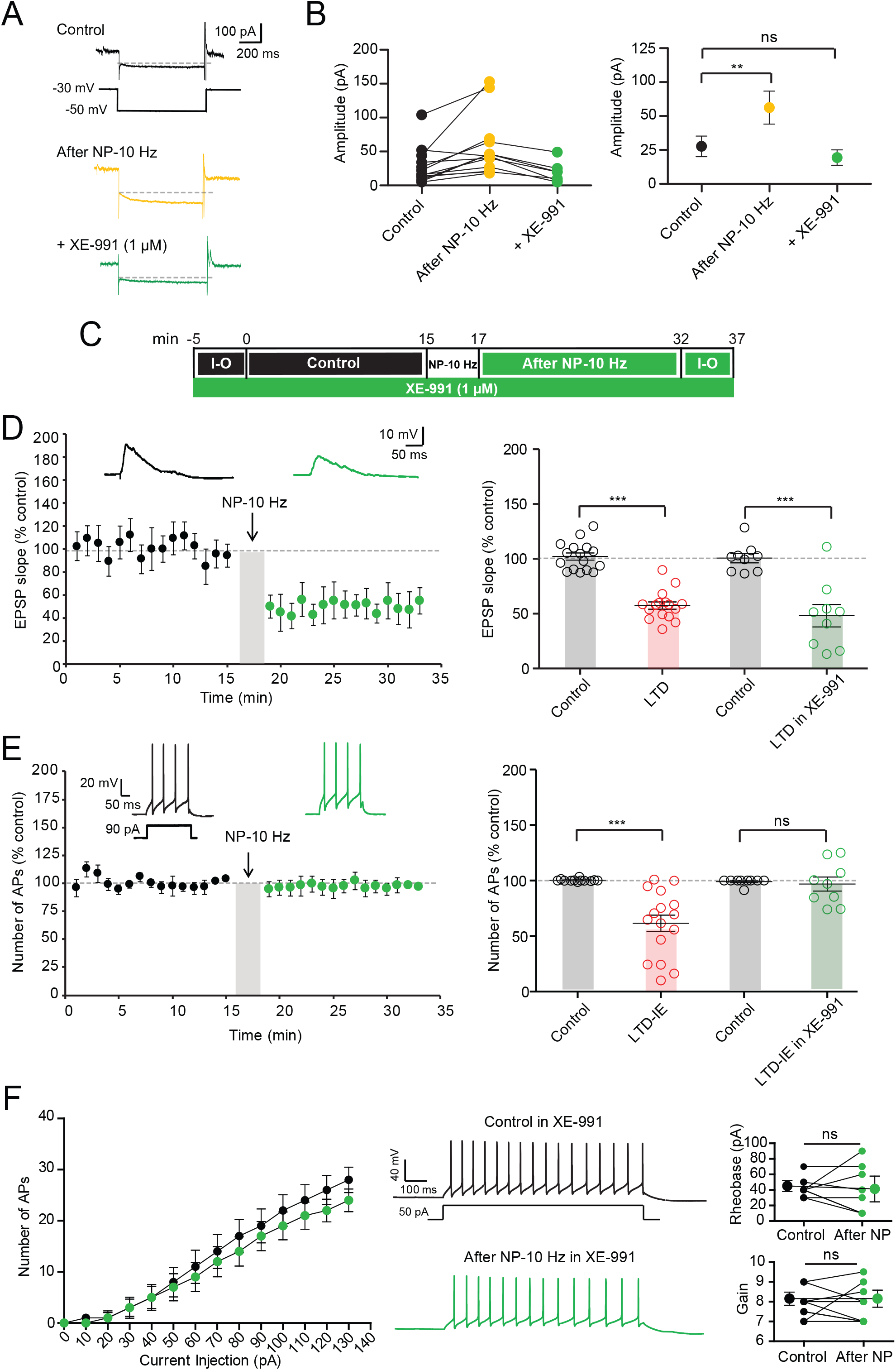
Up-regulation of Kv7 channel-mediated current is involved in LTD-IE expression. A. Up-regulation of M-type current following induction of LTD-IE. O-LM interneurons are recorded in voltage-clamp and the relaxing component of the M-type current is measured in the presence of HCN channel blocker ZD-7288 (1 μM) before (black) and after (yellow) negative pairing, and in the presence of XE-991 (green). B. Pooled data. C. Experimental protocol for LTD and LTD-IE induction in the presence of the Kv7 antagonist, XE-991 (1 μM). D. LTD is induced in the presence of XE-991. E. LTD-IE is totally absent in the presence of XE-991. F. Input-output curves showing the lack of change in both rheobase and gain.

Since M-current is up-regulated, we next checked whether LTD-IE was specifically occluded in the presence of XE-991. As expected, LTD-IE (97 ± 6%, n = 9) but not LTD (49 ± 10%, n = 9) was found to be blocked in the presence of 1 μM XE-991 (**Fig. 3C-E**). In addition, no difference in the rheobase nor in the gain was observed on the input-output curves (**Fig. 3F**). Altogether, these results show that Kv7 channels in O-LM interneurons are up-regulated following induction of LTD by negative pairing induced at theta frequency.

### eCB synthesis mediates both LTD and LTD-IE in O-LM interneurons

Induction of LTD in O-LM interneurons depends on stimulation of the eCB receptor CB1 (18). We first checked whether LTD and LTD-IE were both mediated by CB1 receptors. In the presence of the selective antagonist of CB1 receptor, AM-251 (2 μM), no LTD was induced (95 ± 6%, n = 11; Wilcoxon, p>0.1) but LTD-IE was virtually unchanged (82 ± 5%, n = 11; **Fig. S4**). Furthermore, the rheobase was significantly elevated following negative pairing (from 34.0 ± 5.0 pA to 71.0 ± 12.4 pA, n = 11; Wilcoxson, p<0.01; **Fig. S4**). These results suggest that CB1 receptors mediate LTD of synaptic excitation in O-LM interneurons via a presynaptic modulation of glutamate release but that LTD-IE does not depend on CB1.

Kv7 channels expressed in oocytes are up-regulated by eCBs (10). As, Kv7 channels are involved in LTD-IE, we therefore tested whether inhibiting eCB synthesis prevented LTD-IE. 2-arachidonoylglycerol (2-AG) is an eCB synthetized by diacylglycerol (DAG) lipase. We therefore tested whether the inhibitor of DAG lipase, tetrahydrolipstatin (THL, also called orlistat), blocked induction of LTD-IE. Application of THL (10 μM) for at least 20 minutes was found to slightly increase O-LM excitability (**Fig. 4A**). In fact, the rheobase was significantly diminished in the presence of THL (44.6 ± 4.5 pA, n = 12 in control solution vs. 24.6 ± 4.1 pA, n = 12 in THL), suggesting that basal synthesis of 2-AG is sufficient to reduce excitability in O-LM cells. But most importantly, both LTD and LTD-IE were found to be prevented by inhibiting 2-AG synthesis with THL (**Fig. 4B-D**). In fact, no synaptic modification (99 ± 4% of the control EPSP, n = 12) nor excitability change (100 ± 3% of the control AP number, n = 12) was observed after negative pairing in THL. In addition, no change in the rheobase was observed after negative pairing in THL (**Fig. 4E**). Thus, these results indicate that 2-AG could be a major effector of both synaptic and intrinsic changes in O-LM interneurons.

**Figure 4.**
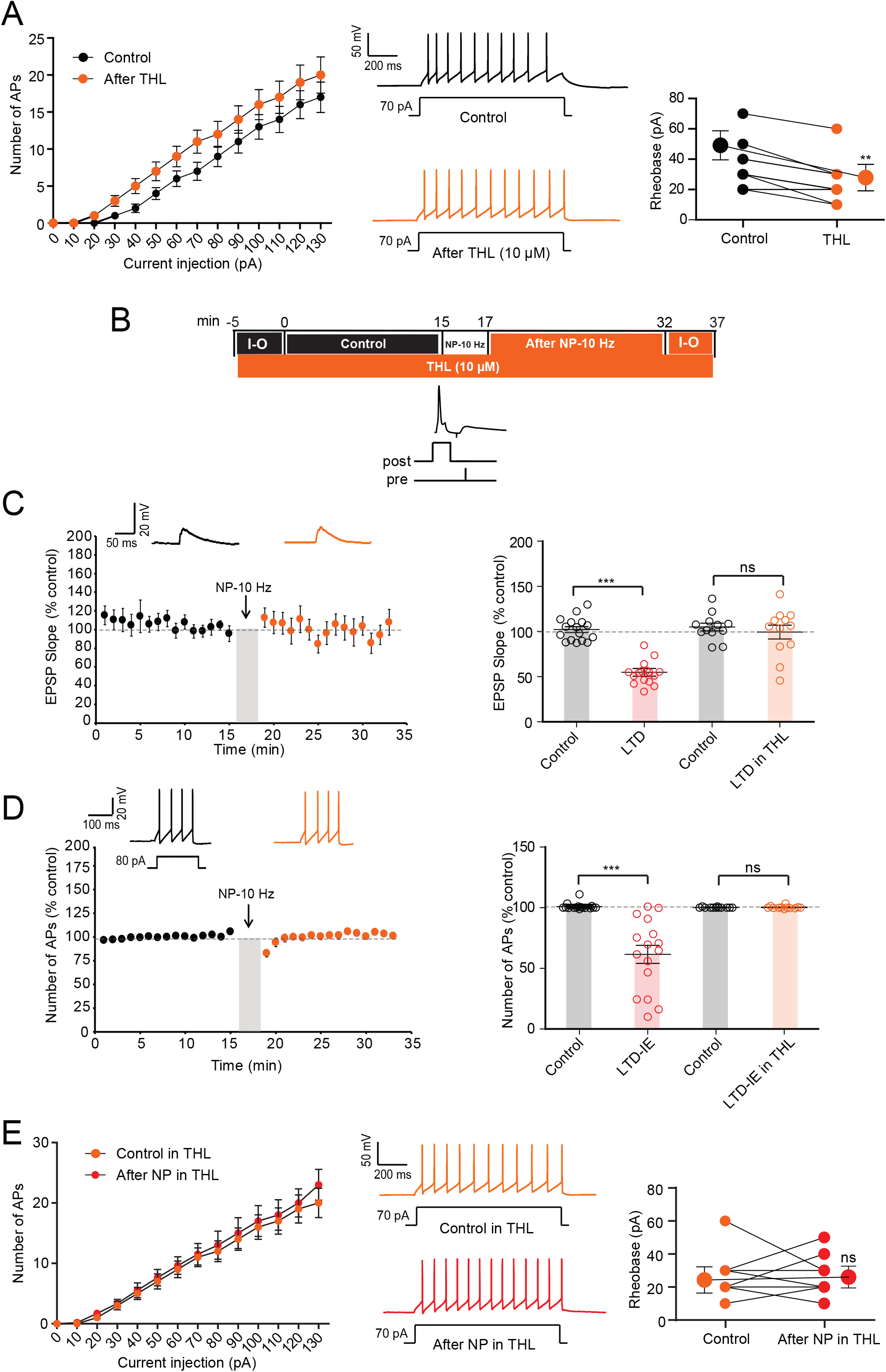
Inhibiting synthesis of 2-AG prevents both LTD and LTD-IE induction. A. Bath application of THL (10 μM) elevates excitability and reduces the rheobase. Left, input-output curves. Middle, discharge before and after application of THL. Right, analysis of rheobase. **, Wilcoxson, p<0.01. B. Experimental protocol for induction of LTD and LTD-IE in the presence of the inhibitor of 2-AG synthesis, THL (10 μM). C. In the presence of THL, no LTD is induced. D. Lack of LTD-IE induction in the presence of THL. E. Input-output curves showing the lack of change in excitability and rheobase. Left, input-output curves. Middle, discharge before and after application of THL. Right, analysis of rheobase.

As eCB synthesis inhibition increased excitability, we next verified that application of eCBs reduced intrinsic neuronal excitability in O-LM interneurons. As expected, application of 2-AG (30 μM) reduced O-LM excitability (rheobase in control: 31.4 ± 2.6 pA, n = 7; rheobase in 2-AG: 45.7 ± 4.3 pA, n = 7; Wilcoxson, p<0.05; **Fig. 5A**). Similarly, JZL-184, an inhibitor of the monoacylglycerol lipase (MAGL), that leads to elevation of 2-AG levels, also reduced excitability through an elevation of the rheobase (from 35.0 ± 4.7 pA, n = 12 to 47.5 ± 5.2 pA; Wilcoxson, p<0.01; **Fig. S5**). Furthermore, application of arachidonoyl-L-serine (ARA-S), a potent activator of Kv7 channels found in the brain (10, 25), reduced O-LM excitability (rheobase in control: 33.7 ± 3.2 pA, n = 8; rheobase in ARA-S: 50.0 ± 5.7 pA, n = 8; Wilcoxson test, p<0.05; **Fig. S5**). In all cases, this reduced excitability was associated with a reduction of the input resistance tested with subthreshold depolarizing pulses (2-AG: from 500 ± 49.2 MΩ to 314.5 ± 23.9 MΩ, n = 7; Wilcoxson test, p<0.05; **Fig. 5B**; JZL-184: from 560.9 ± 52.44 MΩ to 417.3 ± 46.1 MΩ, n=8; Wilcoxson test, p<0.005; **Fig. S5C**; ARA-S: from 545 ± 47.2 MΩ to 406 ± 62.55 MΩ, n = 8; **Fig. S5D**; Wilcoxson test, p<0.05), indicating that this reduced excitability is mediated by M-type current. In the presence of 2-AG, LTD-IE was occluded (**Fig. 5C-D**) and no change in rheobase was observed (rheobase in 2-AG: 45.7 ± 4.2 pA, n = 7; rheobase in 2-AG after NP: 58.5 ± 3.4 pA, n = 7; Wilcoxson, p>0.05; **Fig. 5E**). Similarly, no change in rheobase was observed after negative pairing performed in the presence of JZL-184 or ARA-S (rheobase in JZL-184: 47.5 ± 5.2 pA, n = 7; rheobase in JZL-184 after NP: 50.0 ± 4.7 pA, n = 7; Wilcoxson, p>0.05; **Fig. S5E**; rheobase in ARA-S: 50.0 ± 5.7 pA, n = 8; rheobase in ARA-S after NP: 53.8 ± 5.9 pA, n = 8; Wilcoxson, p>0.05; **Fig. S5F**), indicating that eCBs not only account for induction of LTD but also for expression of LTD-IE in O-LM interneurons.

**Figure 5.**
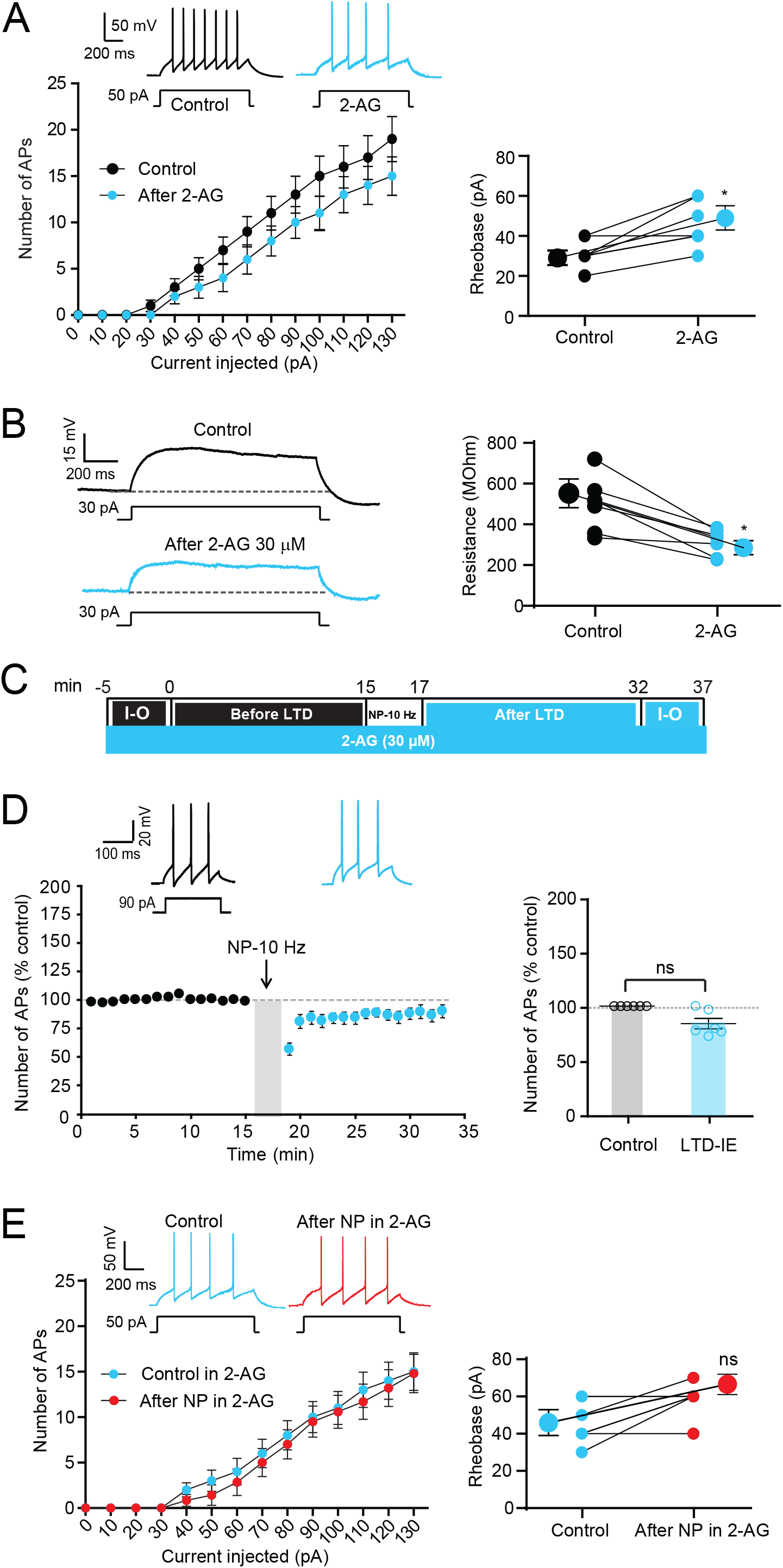
Application of 2-AG mimics and occludes LTD-IE A. Bath application of 2-AG (30 μM) reduces intrinsic excitability and elevates the rheobase in O-LM interneurons. Left, input-output curves and discharge before and after application of 2-AG. Right, analysis of rheobase. *, Wilcoxson, p<0.05. B. 2-AG reduces input resistance tested with depolarizing subthreshold current pulses. Left, representative traces. Right, pooled data. *, Wilcoxson, p<0.05. C. Experimental protocol for induction of LTD-IE in the presence of 2-AG. D. 2-AG occludes induction of LTD-IE. Left, time course. Right, pooled data. E. Input-output curves showing the lack of change in excitability and rheobase. Left, input-output curves and discharge before and after negative pairing in the presence of 2-AG. Right, analysis of rheobase.

### Molecular interaction of eCBs with Kv7.2/3 channels

To confirm the direct molecular interaction of eCBs with Kv7.2/3 channels, we used a modelling approach. Viewed above the membrane, the channel shows its typical homotetramer topology, with the pore located at the center of the structure (**Fig. 6A**). The docking of 2-AG on the 3D structure of rat Kv7.2 in the apo state reveals a high affinity binding site for 2-AG located at the junction of subunits *B* and *C* in the outer leaflet of the plasma membrane (**Fig. 6A** and **6B**). This 2-AG binding site is ideally located to affect the conformational state of the channel. Given the tetrameric structure of the Kv7.2 channel, three other molecules of 2-AG could bind simultaneously to the channel in a symmetric manner, giving a 2-AG stoichiometry of 4 molecules per Kv7.2 channel. 2-AG showed a curved shape that fits well with the binding pocket at the junction of subunits *B* and *C* (**Fig. 6B**). This binding site consists of a gutter formed by amino acid residues 198-209 in chain *B* and 245-252 in chain *C* (**Table S1**; **Fig. 6C** and **6D**).

**Figure 6.**
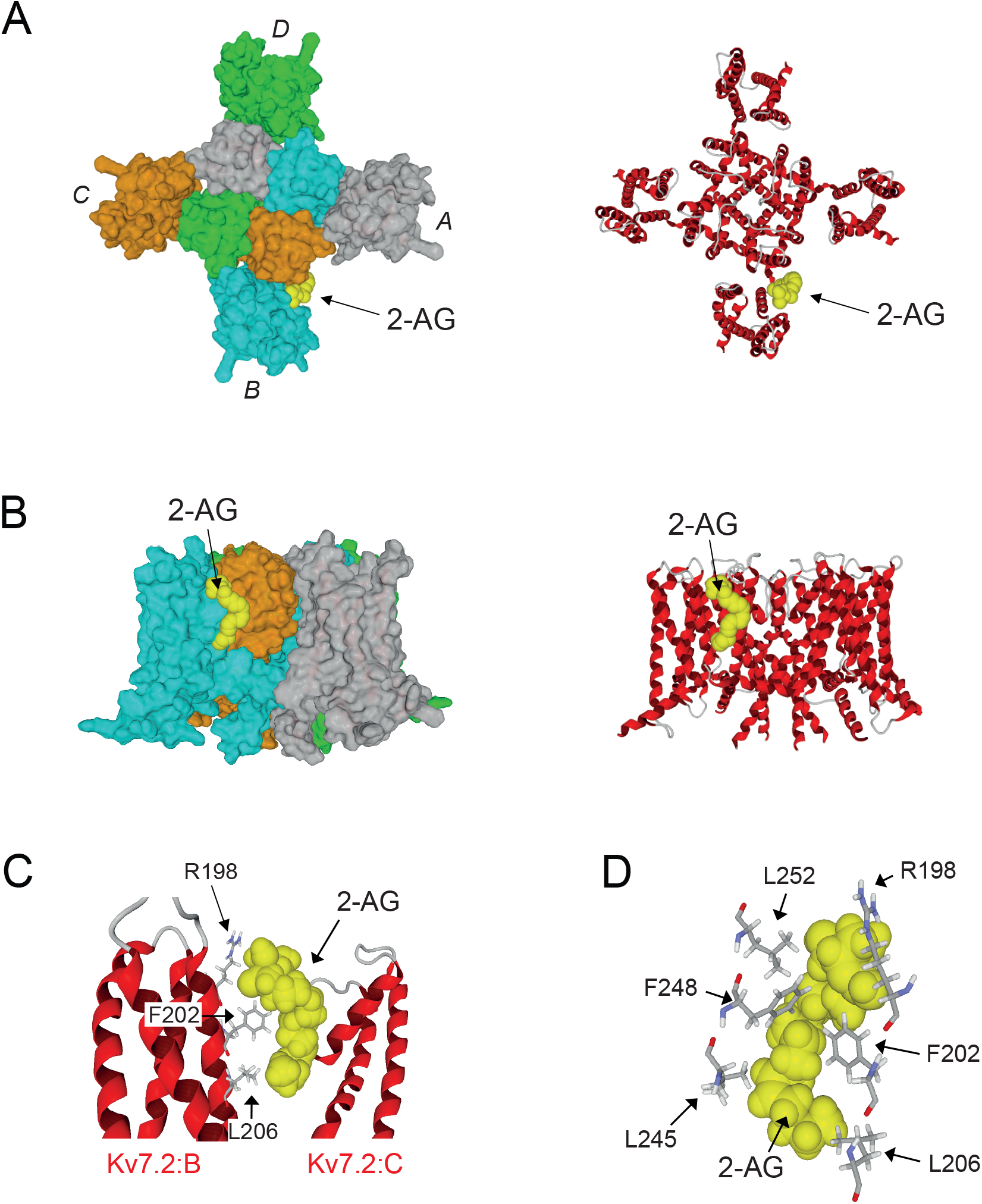
Docking and binding of 2-AG on Kv7.2 in the apo state. A. Kv7.2 consists of a homotetramer of 4 subunits (*A*, *B*, *C* and *D*, respectively colored in grey, cyan, orange and green in the left panel). The pore (still closed in the apo state) is located at the center of the structure. The 2-AG binding site is located in the outer leaflet of the plasma membrane at the junction on chains *B* and *C*. The secondary structure of the channel is shown in the right panel. B. Lateral views of the Kv7.2 channel showing the 2-AG binding site. The surface view in the right panel illustrates the involvement of both chains *B* (cyan) and *C* (orange) in the binding site. C. 2-AG in the endocannabinoid binding pocket formed by subunits *B* and *C* of Kv7.2. Contacts with key amino acid residues of the CARC domain (Arg-198, Phe-202 and Leu-206 in subunit B) are clearly shown. D. Amino acid residues of subunits *B* (Arg-198, Phe-202 and Leu-206) and *C* (Leu-245, Phe-248 and Leu-252) in close contact with 2-AG. For clarity only the key triad of amino acids in each subunit are shown.

ARA-S is a structural analogue of 2-AG and binds to Kv7 (10). The model shows that ARA-S and 2-AG share the same the binding pocket (**Fig. S6A** and **S6B**). The superposition of 2-AG and ARA-S further demonstrates that both eCBs share the same binding site (**Fig. S6C**). Although both eCBs have a high affinity for Kv7.2, the energy of interaction is higher for ARA-S compared with 2-AG (53.6 vs. 36.8 kJ.mol^−1^; **Table S1**). This difference is essentially due to the negatively charged carboxylate in the polar head group of ARA-S that forms an electrostatic bond with the cationic group of Arg-198 and to the van der Waals contact with Leu-206 which is closer to ARA-S than to 2-AG. Both eCBs adopt the same typical curved structure that fits with the binding site. It is interesting to note that the eCB binding site is highly conserved in Kv7.2 and Kv7.3. The amino acid sequence of the binding motif in chain *B* (198-209) is identical for both channels: RSLRFLQILRMI.

The presence of a CARC motif within the eCB binding domain suggests that cholesterol could also bind to Kv7.2. Docking studies confirmed this hypothesis **(Fig. S6D)**. The same amino acid residues that interact with 2-AG (i.e. Arg-198, Phe-202, and Leu-206; **Fig. S6D**) are also involved in cholesterol recognition (**Fig. S6D**). Interestingly, when 2-AG was docked onto the Kv7.2 conformer pre-constrained by cholesterol, the binding of the eCB was improved, due to the reorientation of aromatic rings (residues Phe-202 and Phe-248), and an optimized geometry of the side chain of Arg-198 (**Fig. S6D**). Overall, the energy of interaction of 2-AG was raised from −36.8 to − 53.5 kJ.mol^−1^ (**Table S2**). Thus, although our data suggest that eCBs can bind to Kv7.2 in a cholesterol-independent mechanism, it is likely that cholesterol could further improve the binding of 2-AG by a conformational selection process. The higher flexibility of 2-AG compared to the sterol, together with the higher energy of interaction of the Kv7.2 complex (**Table S2**) is compatible with a displacement of cholesterol by 2-AG, leading to the formation of a stable, long lasting eCB-Kv7.2 complex. We conclude that this eCB-Kv7 complex may account for the reduced excitability found in O-LM interneurons.

## DISCUSSION

Our study reveals that O-LM interneurons unexpectedly express intrinsic plasticity in parallel to synaptic plasticity. In fact, we show here that negative correlation between pre- and post-synaptic activity in the theta range induces long-lasting depression of both synaptic transmission and intrinsic neuronal excitability. These two modifications are synergistic and provide a certain level of functional redundancy because both types of plasticity tend to reduce the excitation of O-LM interneurons. Mechanistically, our study indicates that both LTD and LTD-IE in O-LM interneurons depend on the biosynthesis of eCBs that target 1) presynaptic CB1 receptors to regulate release of glutamate and 2) post-synaptic Kv7 channels present in the dendrites of O-LM interneurons to regulate intrinsic excitability. This mechanism of LTD-IE can potentially have very important consequences in the feedback loop that O-LM interneurons establish with CA1 pyramidal neurons in the local hippocampal circuit.

### Functional synergy between synaptic and intrinsic modifications

Long-lasting synaptic plasticity is generally associated with synergistic regulation of intrinsic neuronal excitability (21). For instance, in hippocampal pyramidal neurons, both LTP and LTD are respectively induced in parallel with E-S potentiation and E-S depression (26, 27). In fast spiking PV interneurons, both LTP and LTP-IE are induced by high frequency stimulation of glutamatergic inputs (22). The present study shows that this rule is also verified in O-LM interneurons since LTD and LTD-IE occur in parallel. Interestingly, a positive correlation was found between LTD and LTD-IE values. In addition, pre- or post-synaptic activation alone was found to be unable to induce both LTD and LTD-IE, suggesting that induction of both LTD and LTD-IE are mechanistically linked.

Our study points to the conclusion that eCBs play a critical role in induction of both LTD and LTD-IE in O-LM interneurons. First, inhibition of CB1 receptor with AM-251 prevented LTD induction. Second, inhibition of DAG lipase by THL totally abolished both LTD and LTD-IE induction. Third, application of eCBs as 2-AG or ARA-S reduced intrinsic excitability in O-LM interneurons.

### Intrinsic plasticity in GABAergic interneurons

Activity-dependent plasticity has been reported in only a few type of cortical interneurons including peri-somatic inhibitory interneurons such as PV baskets cells (22, 23) or GABAergic interneuron-targeting neurogliaform interneurons (28). O-LM interneurons thus represent the third class of GABAergic interneurons showing plasticity of intrinsic neuronal excitability, suggesting that plasticity of inhibitory circuits is not only mediated by synaptic plasticity but that intrinsic changes take a large part in this regulation. In all three interneuron types, intrinsic plasticity involves the regulation of Kv channels: Kv1.1 for PV basket cells (22), Kv4 for neurogliaform interneurons (28) and Kv7 for O-LM cells (this study). However, while the two other plasticity mechanisms involve a loss of function, the present plasticity reveals a gain of function.

### Involvement of Kv7 channels in intrinsic plasticity

We show here that Kv7 channels represent the main ion channel by which neuronal excitability in O-LM interneurons is regulated by synaptic activity. The M-current was found to be of greater amplitude after negative pairing. Furthermore, LTD-IE expression (but not LTD) was prevented in the presence of the selective antagonist of Kv7 channels, XE-991. Although Kv7 channels have been suspected to be involved in functional plasticity (29), our study provides the first direct evidence for its involvement in activity-dependent intrinsic regulation of intrinsic excitability.

In contrast to most neurons, the dendrites of O-LM interneurons contain a high density of Nav channels that makes possible AP initiation in the dendrites (30). Therefore, Kv7 channels located in the dendrites (7) may critically control O-LM excitability. Kv7 channels are also present in CA3 basket cells and in bistratified interneurons (31). Interestingly, PV basket interneurons express cannabinoid-dependent LTD (18), suggesting that cannabinoid-dependent LTD-IE could be induced in these interneurons.

### eCB signaling in O-LM interneurons

eCBs (especially 2-AG) have emerged as very important retrograde messenger throughout the brain. eCBs are involved in many brain functions including regulation of presynaptic transmitter release underlying learning and memory (32), neurological disorders (33) and psychiatric disorders (34). Interneurons function is also controlled by eCB. O-LM interneurons express high levels of the DAG lipase-α in perisynaptic regions around excitatory inputs (35, 36). In addition, a specific eCB-dependent presynaptic LTD has been described in O-LM cells (18).

Our results show that 2-AG or ARA-S application reduces intrinsic excitability of O-LM interneurons. In fact, both compounds elevate the rheobase and partially occlude induction of LTD-IE. A recent study indicates that ARA-S facilitates activation of the M-current mediated by Kv7.2/3 in an *in vitro* expression system (10). In fact, ARA-S hyperpolarizes the activation curve by and increases by a factor 2 the conductance of the M- current. However, in that study, no indication about a potential persistent effect of ARA-S was identified. Our molecular modelling indicates that 2-AG or ARA-S bind to Kv7.2 at the junction between two adjacent subunits with a very high affinity, suggesting a persistent binding and possibly a persistent activation of these channels. The identified eCB binding site is highly conserved in Kv7.2 and Kv7.3. The amino acid sequence of the binding motif in chain B (198-209) is identical for both channels: 198-<u>**R**</u>SLR<u>**F**</u>LQI<u>**L**</u>RMI-209. This motif is a consensus CARC domain, with the key amino acid residues bold and underscored. Such CARC motifs in TM domains are recognized by cholesterol (37). Thus, our data suggest the intriguing possibility of a regulatory cholesterol-eCB exchange in Kv7 channels function, in line with previous studies focused on CB1 receptors (38). The binding motif in chain *C* of Kv7.2 has the following amino acid sequence in Kv7.2: 245-LASFLVYL-252, which belongs to an inverted CARC motif (37) that encompasses residues 245-255 (<u>**L**</u>ASFLV<u>**Y**</u>LAE<u>**K**</u>). This motif is also conserved in Kv7.3 (<u>**L**</u>SSFLV<u>**Y**</u>LVE<u>**K**</u>). Taken together, these data further strengthen the hypothesis of a dual cholesterol-eCB regulation of Kv7.2 and Kv7.3 function.

Our data indicate that the regulation of Kv7 channels by eCBs is not mediated by CB1 receptor but rather, eCBs may directly interact with Kv7 channels. First, inhibiting CB1 receptors with AM-251 blocked induction of LTD but not LTD-IE. Furthermore, molecular modeling indicates that 2-AG is able to directly bind to Kv7.2/3 channels.

Lipids as eCBs are known to control the biophysical properties of a wide range of ion channels (39), including Kv1 (40, 41), Kv4 (42) and HCN (43) channels that are involved in long-term plasticity of intrinsic excitability (22, 44–47). Thus, our study may lead to open a new line of future investigations.

### Functional consequences

The functional consequences of a reduction in O-LM excitability mediated by the up-regulation of Kv7 channel-mediated M-type current may be multiple. At the cellular level, up-regulation of M-type current may enhance intrinsic resonance in O-LM cells in the depolarizing range. In fact, both Kv7 and HCN channels are known to be responsible for intrinsic resonance at two specific voltage levels in CA1 pyramidal neurons (48). O-LM interneurons possess these two channels and also display intrinsic resonance (24). At the local circuit level, a reduced excitability of O-LM interneurons may reduce inhibition at distal dendrites of CA1 pyramidal neurons and thus favor induction of LTP at temporo-amonic pathway (6). In parallel, the reduced inhibition of *stratum radiatum* interneurons may consequently reduce LTP induction at Schaffer collateral pathway (6). This switch can potentially have an important role for the acquisition of new spatial memories and alteration could result in pathological consequences. Furthermore, the eCB-dependent dampening of O-LM excitability may represent a way to limit excitation during excessive network excitation. Further studies will be required to circumvent the functional consequence of LTD-IE in O-LM interneurons.

## METHODS

### Slices preparation

Hippocampal slices (350 mm) were prepared from postnatal day 14–20 Wistar rats. All experiments were carried out according to the European and institutional guidelines for the care and use of laboratory animals (Council Directive 86/609/EEC and French National Research Council) and approved by the local health authority (Préfectture des Bouches-du-Rhône, Marseille). Rats were deeply anesthetized with isoflurane and killed by decapitation. Slices were cut with a vibratome (Leica VT-1000S) in a N-methyl-D-glucamine (NMDG) solution containing (in mM: 92 NMDG, 1.2 NaH_2_PO_4_, 30 NaHCO_3_, 20 Hepes, 25 D-glucose, 5 sodium ascorbate, 2 thiourea, 3 sodium pyruvate, 10 MgCl_2_ and 0.5 CaCl_2_). The slices were maintained for one hour at room temperature in oxygenated (95%O_2_/5%CO_2_) artificial cerebrospinal fluid (ACSF; in mM: NaCl, 125; KCl, 2.5; NaH_2_PO_4_, 0.8; NaHCO_3_, 26; CaCl_2_, 3; MgCl_2_, 2; D-Glucose, 10).

### Electrophysiology

Each slice was transferred to a temperature-controlled (31°) recording chamber with oxygenated ACSF. O-LM hippocampal interneurons were identified by the location of their soma (*stratum oriens* of the CA1), their morphology (a spindle-shaped cell body horizontally oriented along the pyramidal layer) and their peculiar electrophysiological signature (the characteristic “sag” depolarizing potential in response to hyperpolarization currents injections and the typical “saw-tooth” shape of action-potential afterhyperpolarizations). In all recordings GABA_A_ receptors were blocked by picrotoxin (PTx 100 mM) and CA3 area was surgically removed to prevent epileptiform bursting. Whole-cell patch-clamp recordings were obtained from CA1 O-LM interneurons. The electrodes were filled with an internal solution containing (in mM): K-gluconate, 120; KCl, 20; HEPES, 10; EGTA, 0; MgCl_2_6H_2_O, 2; Na_2_ATP, 2. Stimulating pipettes filled with extracellular saline were placed in the stratum oriens near the alveus in order to stimulate inputs coming from the pyramidal neurons.

In control and test conditions the following parameters were measured in current clamp mode for 15 minutes before and at least 15 minutes after LTD protocol: apparent input resistance was tested by current injection (−120 pA; 800 ms); EPSPs were evoked at 0.1 Hz and the stimulus intensity (100 μs, 40–100 μA) was adjusted to evoke subthreshold EPSPs (4–10 mV). Short current injections (70-120 pA, 100 ms) were applied at each sweep in order to measure the intrinsic excitability as the number of spikes over time. Series resistance was monitored throughout the recording and only experiments with stable resistance were kept (changes <10%). Before and after LTD induction, a protocol was designed in order to plot input-output curves by measuring action potentials number in response to incrementing steps of current pulses.

To measure M-current mediated by somato-dendritic Kv7 channels, a voltage clamp protocol was used consisting in voltage steps from −30 mV to −50 mV (Lawrence et al., 2006). The relaxing component of the M current was then measured before and after LTD-IE induction.

### Induction of LTD and LTD-IE

LTD and LTD-IE were induced in O-LM interneurons with a protocol consisting of 600 pairings between a post-synaptic action potential followed by a presynaptic stimulation of the pyramidal neurons with a delay of 10 ms. These pairings were evoked at a frequency of 5, 10 or 20 Hz in blocks of 6 with an inter-block interval of 10 seconds (18). For each cell we used a protocol, which allowed us to measure excitatory post-synaptic potentials (EPSPs), input resistance and intrinsic excitability (IE), 15 minutes before and at least 15 minutes after the LTD protocol.

### Data acquisition and analysis

Recordings were obtained using a MultiClamp 700B (Molecular Devices) amplifier and pClamp10 software. Data were sampled at 10 kHz, filtered at 3 kHz, and digitized by a Digidata1322A (Molecular Devices). All data analyses were performed with custom written software in Igor Pro 6 (Wavemetrics). Apparent input resistance was determined by the subtraction of the steady-state voltage change during hyperpolarizing current injection from the baseline. Pooled data are presented as means ± SEM. Statistical comparisons were made using Mann–Whitney or Wilcoxon tests with Prisma GraphPad software.

### Pharmacology

Drugs were all bath applied. Picrotoxin (PiTx) was purchased from Sigma; [4-(N-ethyl-N-phenylamino)-1,2-dimethyl-6-(methylamino) pyrimidinium chloride] (ZD-7288), tetrahydrolipstatin (THL), 2 Arachidonylglycerol (2-AG), JZL-184, AM 251 (*N*-(Piperidin-1-yl)-5-(4-iodophenyl)-1-(2,4-dichlorophenyl)-4-methyl-1*H*-pyrazole-3-carboxamide) from Tocris and; N-Arachidonoyl-L-Serine (ARA-S) from Cayman Chem.

### Morphological analysis

The dendritic and axonal morphology of the recorded neurons was revealed by biocytin staining. For this, biocytin (0.2−0.4%, Sigma) was added to the pipette solution and was revealed with avidin-biotin complex coupled to fluorescein, and examined using confocal microscopy (Zeiss, LSM-780). The morphology of the neurons was reconstructed using ImageJ.

### Molecular modeling

The 3D structure of the rat Kv7.2 channel has been obtained by homology with the human KCNQ2 protein in apo state (49). The amino acid sequence of rat Kv7.2 (96.61% sequence homology) was retrieved from UniProtKB −O88943 (KCNQ2_RAT) and submitted to the Swiss Model server (SWISS-MODEL Interactive Workspace (expasy.org)). The model is stored in the Swiss Model database (project “KCNQ2_RAT O88943 Potassium voltage-gated channel subfamily KQT member 2”).

Docking of ara-S and 2-AG on Kv7.2 was performed with Hyperchem as previously described (50). Energy minimization of each system was performed with the Polak-Ribière conjugate gradient algorithm, with the Bio-CHARMM force field, using a maximum of 3×10^5^ steps, and a root-mean-square (RMS) gradient of 0.01 kcal. Å^−1^.mol^−1^ as the convergence condition. The energies of interaction were calculated by the Ligand Energy Inspector function of Molegro Molecular viewer (Molexus IVS, Denmark http://molexus.io/molegro-molecular-viewer/).

## Data availability

All study data are included in the article

## Author contribution

SI and DD designed research. SI, MS & MR performed experimental research. JF made the modeling. SI, NA, JF & DD analyzed data, and SI, JF & DD wrote the manuscript that was approved and amended by all authors.

## Acknowledgments

We thank JJ Ramirez-Franco for help with use of the confocal microscope. This work was supported by INSERM, CNRS, Aix-Marseille Université, NeuroMarseille and Fondation pour la Recherche Médicale (FRM DEQ20180839583 to DD).

---

**Figure S1.**
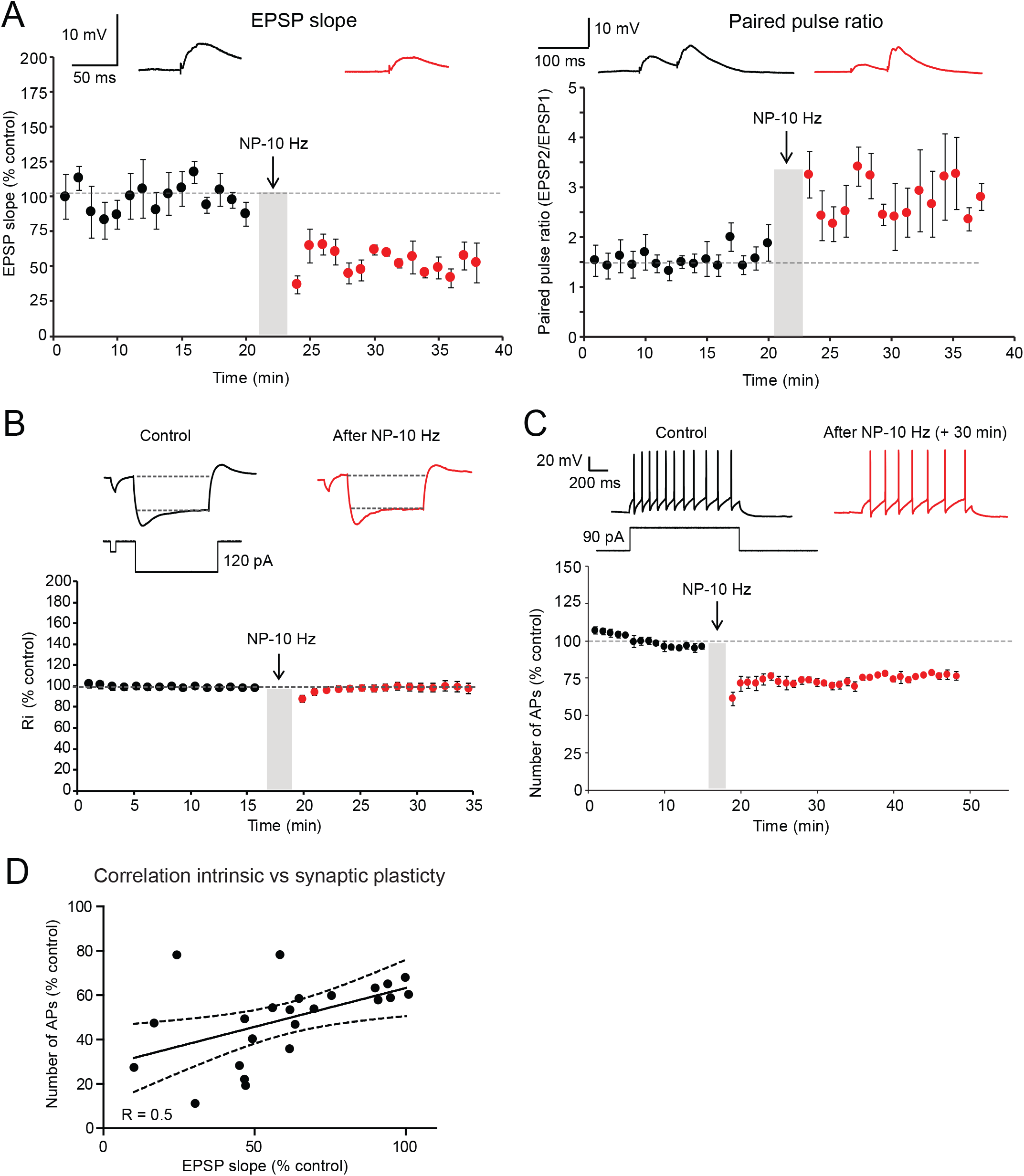
Induction of LTD and LTD-IE in O-LM interneurons. A. LTD in O-LM interneurons is associated with an increased paired-pulse ratio. Left, time-course of synaptic depression induced by negative pairing at 10 Hz. Right, paired-pulse ratio is increased. B. Lack of input resistance change after negative pairing. Up, representative traces. Bottom, time-course of normalized input resistance. C. Long-lasting reduction in intrinsic excitability. The reduction in excitability can be followed during more than 30 minutes. D. Correlation between normalized AP number and normalized EPSP slope.

**Figure S2.**
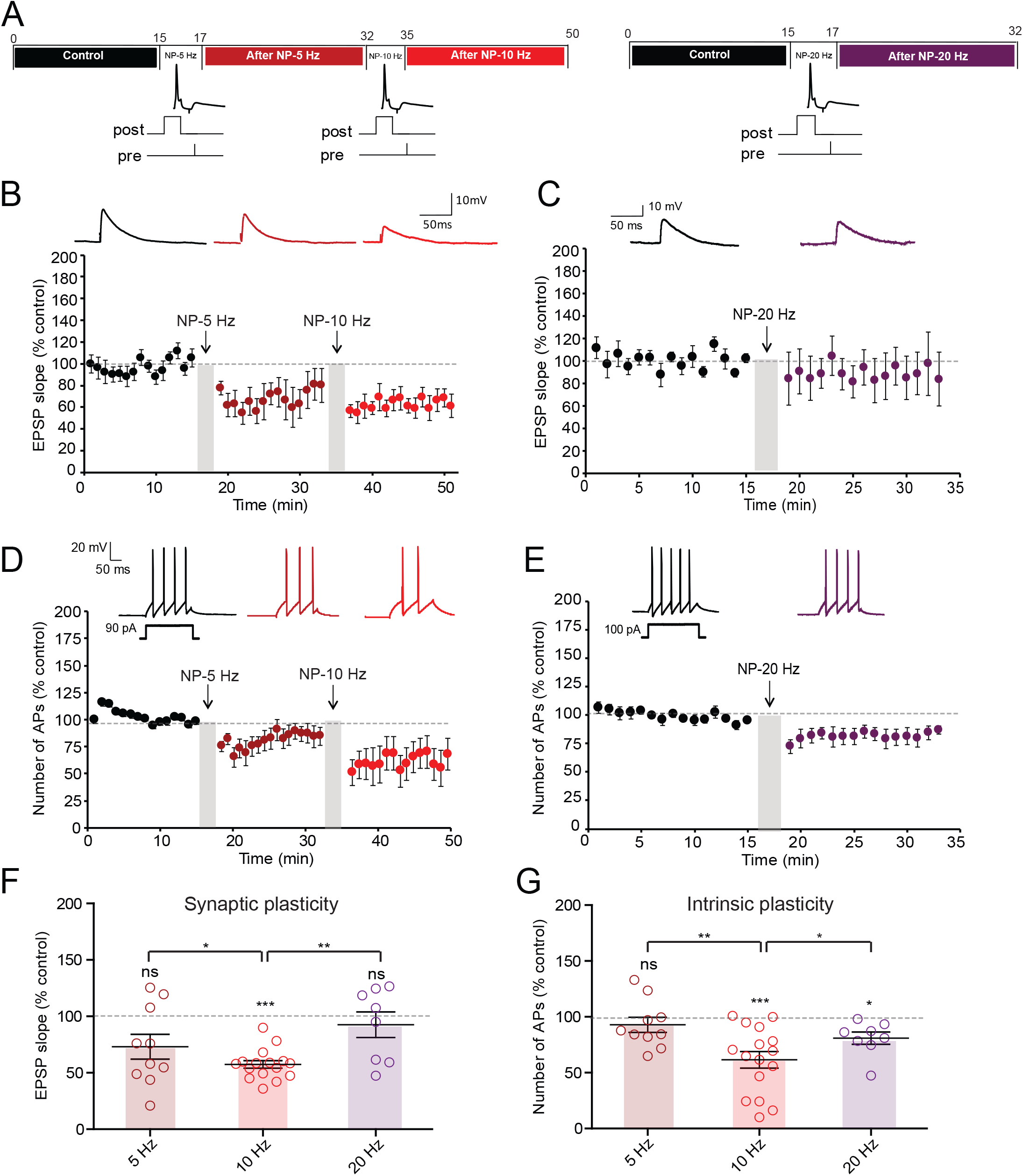
LTD-IE is induced by stimulation at 10 Hz. A. Experimental protocols. Left, negative pairing at 5 Hz is followed by negative pairing at 10 Hz. Right, negative pairing at 20 Hz. B. Synaptic changes after negative pairing at 5 Hz and 10 Hz. C. Synaptic changes following negative pairing at 20 Hz. D. Intrinsic changes following negative pairing at 5 and 10 Hz. E. Intrinsic changes following negative pairing at 20 Hz. F. Summary of synaptic changes as a function of frequency. G. Summary of intrinsic changes as a function of frequency.

**Figure S3.**
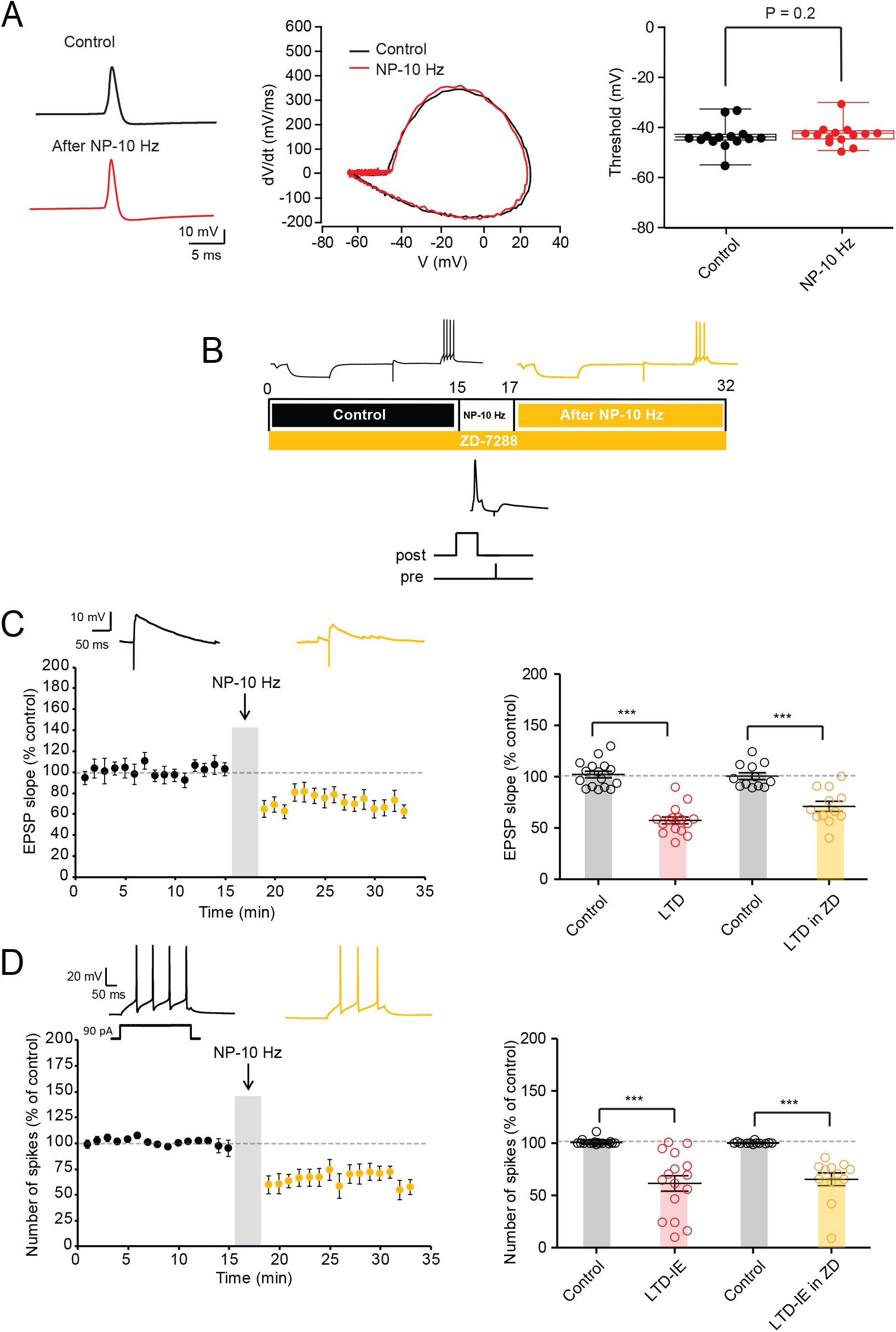
LTD-IE expression: unchanged AP threshold and no incidence of HCN inhibition. A. O-LM AP threshold remains stable following induction of LTD-IE. Left, spike before (black) and after (red) negative pairing. Middle, phase plots of the spike. Right, pooled data. B. Experimental protocol for induction of LTD and LTD-IE in the presence of 1 μM ZD-7288. C. Synaptic changes induced by negative pairing in the presence of ZD-7288. Left, time course. Right, comparison with control condition. D. Intrinsic changes following negative pairing in the presence of ZD-7288. Left, time course. Right, comparison with control condition.

**Figure S4.**
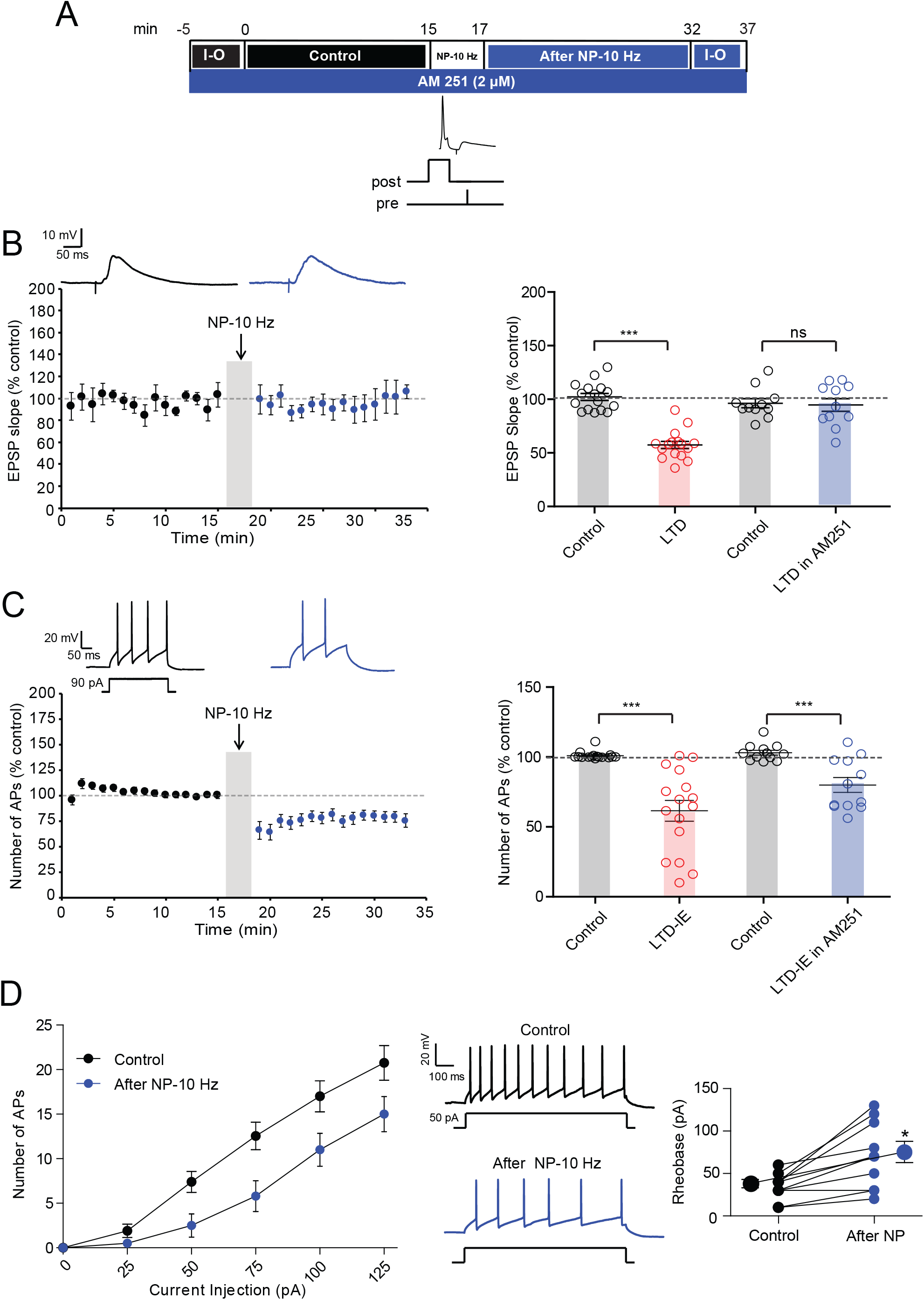
Inhibition of CB1 receptor block LTD but not LTD-IE. A. Experimental protocol for induction of LTD and LTD-IE in the presence of 2 μM AM-251. B. Synaptic changes induced by negative pairing in the presence of AM-251. Left, time course showing a lack of LTD. Right, comparison with control condition. C. Intrinsic changes following negative pairing in the presence of AM-251. Left, time course. Right, comparison with control condition. D. Input-output curves obtained in control and following negative pairing in the presence of AM-251. Left, curves. Middle, discharge before and after negative pairing. Right, analysis of rheobase.

**Figure S5.**
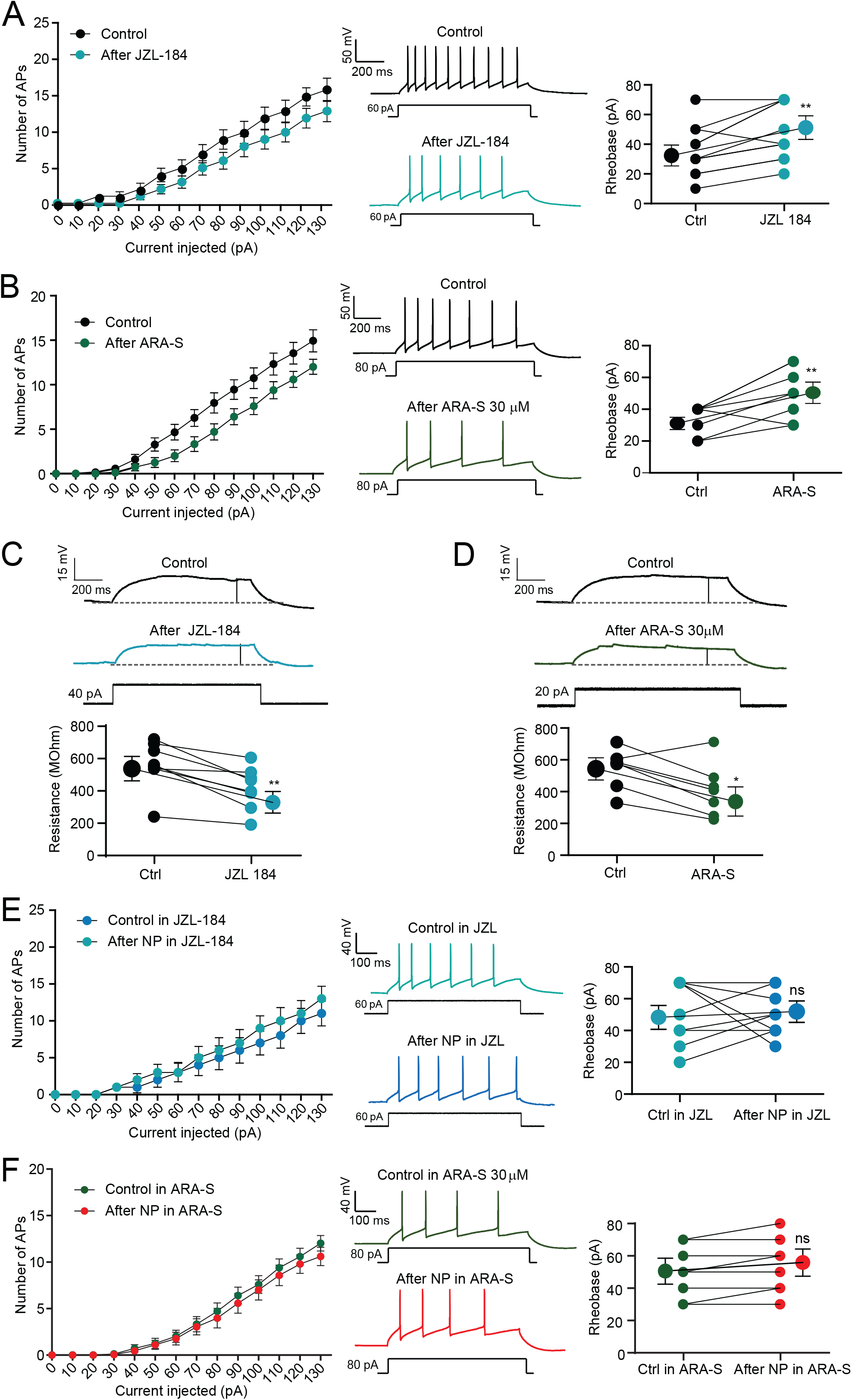
Effects of endocannabinoids on O-LM excitability. A. The 2-AG degradation inhibitor, JZL-184 (1 μM) reduces intrinsic excitability in O-LM interneurons. Left, input-output curves. Middle, discharge before and after application of JZL-184. Right, analysis of rheobase. **, Wilcoxson, p<0.01. B. ARA-S (1 μM) reduces O-LM excitability. Left, input-output curves. Middle, discharge before and after application of ARA-S. Right, analysis of rheobase. **, Wilcoxson, p<0.01. C. JZL-184 reduces input resistance tested with subthreshold depolarizing pulses. Top, representative traces. Bottom, pooled data. **, Wilcoxson, p<0.01. D. ARA-S reduces input resistance tested with subthreshold depolarizing pulses. Top, representative traces. Bottom, pooled data. **, Wilcoxson, p<0.01. E. Lack of intrinsic plasticity in the presence of JZL-184. Left, input-output curves. Middle, discharge before and after application of JZL-184. Right, analysis of rheobase. F. Lack of intrinsic plasticity in the presence of ARA-S. Left, input-output curves. Middle, discharge before and after application of ARA-S. Right, analysis of the rheobase.

**Figure S6.**
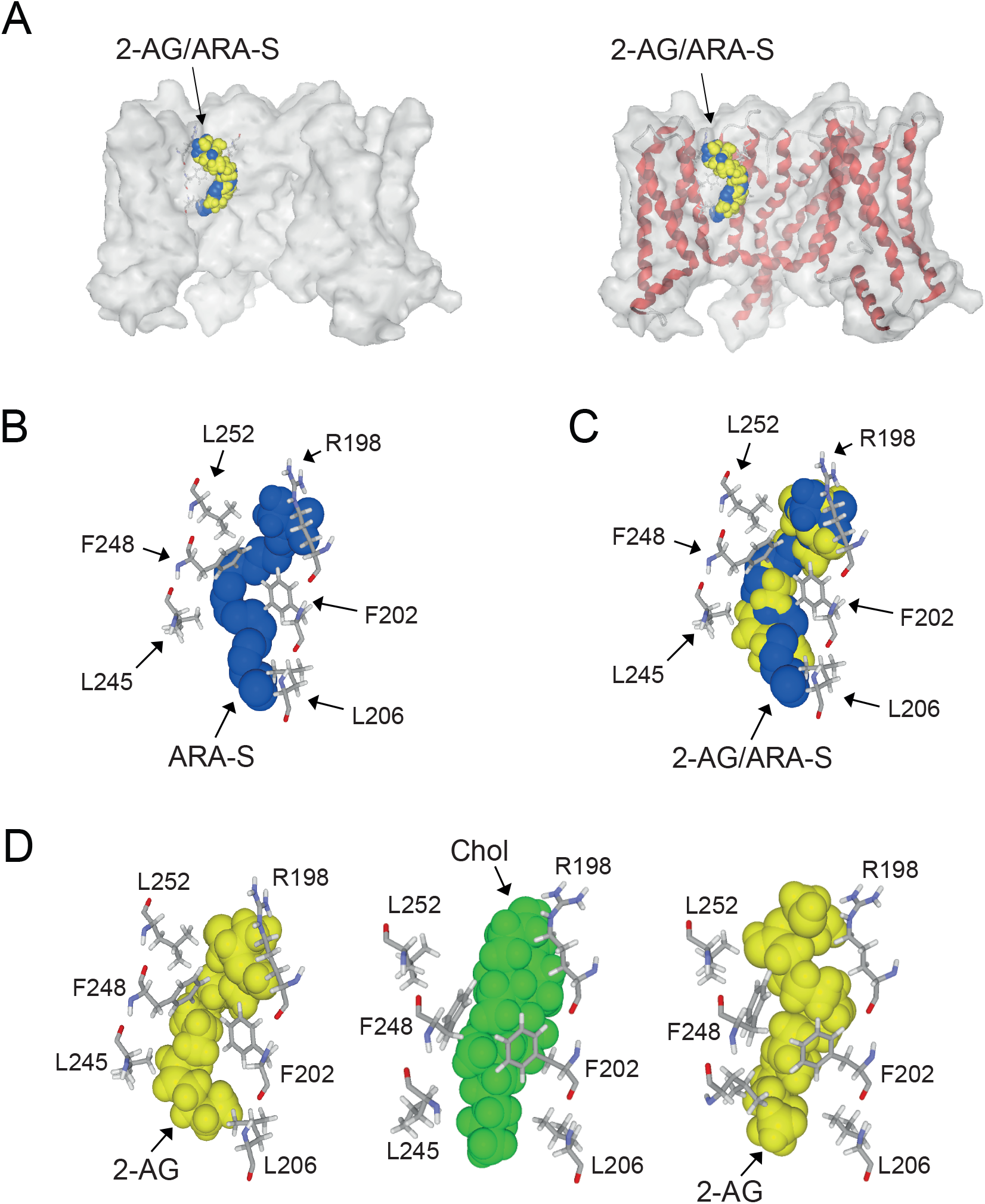
2-AG and ARA-S binding to Kv7.2 A. Superposition of 2-AG (yellow) and ARA-S (blue) bound to Kv7.2 in surface view (left panel) and secondary structure/surface rendition (right panel). Both endocannabinoids occupy a similar volume and bind to the same site. B. Amino acid residues of Kv7.2 in close contact with ARA-S. C. Superposition of 2-AG and ARA-S in the cannabinoid binding pocket of Kv7.2. D. Previous docking of cholesterol enhances interaction of 2-AG for Kv7.2. Left, docking of 2-AG (in yellow) on Kv7.2. Middle, docking of cholesterol (in green) on Kv.2 re-orientates the aromatic rings of Phe-202 and Phe-258 and re-arranges the side chain of Arg-198. Right, docking of 2-AG (in yellow) on the cholesterol-optimized conformation of Kv7.2 shown in the middle. Note the reorientation of Leu-245 which does not interact with cholesterol (middle). The aromatic rings of Phe-202 and Phe-248 have retained the specific orientations imposed by cholesterol.

**Table S1.** Energy of interaction (in kJ.mol^−1^) of each amino acid residue of Kv7.2 involved in 2-AG and ARA-S binding.

| **Amino acid** | **2-AG** | **ARA-S** |
| --- | --- | --- |
| Arg-198 (chain B) | -6.0 | -13.6 |
| Phe-202 (chain B) | -9.4 | -9.5 |
| Leu-206 (chain B) | -2.0 | -7.9 |
| Ile-209 (chain B) | -1.2 | -5.8 |
| Leu-245 (chain C) | -3.5 | -1.7 |
| Phe-248 (chain C) | -8.9 | -9.8 |
| Leu-249 (chain C) | -1.6 | -2.6 |
| Leu-252 (chain C) | -4.2 | -2.7 |
| Total | -36.8 | -53.6 |

**Table S2.** Energy of interaction (in kJ.mol^−1^) of each amino acid residue of Kv7.2 involved in 2-AG binding in absence of cholesterol or after cholesterol release. For comparison, the energy of interaction of cholesterol bound to Kv7.2 is also given.

| **Amino acid** | **2-AG bound**<br>**without cholesterol** | **2-AG bound**<br>**after cholesterol**<br>**release** | **Cholesterol**<br>**bound** |
| --- | --- | --- | --- |
| Arg-198 (chain B) | -6.0 | -14.9 | -13.1 |
| Ser-199(chain B) | 0 | -2.8 | - 3.1 |
| Phe-202 (chain B) | -9.4 | -20.0 | -22.6 |
| Leu-206 (chain B) | -2.0 | -2.5 | -2.4 |
| Ile-209 (chain B) | -1.2 | -1.4 | -0.8 |
| Leu-245 (chain C) | -3.5 | -2.0 | 0 |
| Phe-248 (chain C) | -8.9 | -7.7 | -7.7 |
| Leu-249 (chain C) | -1.6 | 0 | 0 |
| Leu-252 (chain C) | -4.2 | -2.2 | 0 |
| Total | -36.8 | -53.5 | -49.7 |

